# Machine-Learning Based Optimisation of a Biomimiced Herringbone Microstructure for Superior Aerodynamic Performance

**DOI:** 10.1101/2022.09.17.508361

**Authors:** Rushil Samir Patel, Harshal D. Akolekar

## Abstract

Biomimicry involves taking inspiration from existing designs in nature to generate new and efficient systems. The feathers of birds which form a characteristic herringbone riblet shape are known to effectively reduce drag. This paper aims to optimise the individual constituent structure of a herringbone riblet pattern using a combination of computational fluid dynamics (CFD) and supervised machine learning algorithms to achieve the best possible reduction in drag. Initially, a herringbone riblet design is made by computer aided designing and is parameterised. By randomly varying these parameters, 107 additional designs are made and are subjected to CFD calculations to derive their drag coefficients (*C*_*d*_). These designs are used to train a supervised learning model which is employed as an alternative to CFD for predicting the *C*_*d*_ of other 10000 randomly generated herringbone riblet designs. Amongst these, the design with the least predicted Cd is considered as the optimised design. The *C*_*d*_ prediction for the optimised design had an error of 4 % with respect to its true *C*_*d*_ which was calculated by using CFD. The optimised design of this microstructure can be utilised for drag reduction of aeronautical, automotive or oceanic crafts by integrating them onto their surfaces.

## 1. Introduction

The design of an aircraft is aimed at reducing the net specific fuel consumption and increasing the flight efficiency. Different factors such as flight routing, propulsive efficiency and aeroacoustic behaviour can affect the flight performance. Amongst these factors, the fluidic drag on an aircraft is of great importance in affecting the propulsive efficiency. Any object moving through a fluid medium encounters a resistive force which is known as drag. Drag is classified into skin friction drag, lift drag, pressure drag and compressibility drag [1]. The most significant types of drag forces encountered by a cruising commercial aircraft are the skin friction or viscous drag and the drag due to lift or vortex drag [1]. It is estimated that a reduction of 10 % in the viscous drag on an aircraft fuselage can lead to fuel savings of up to $ 1 billion per year for airlines [2]. Therefore, it becomes necessary to reduce the drag experienced by any moving craft. Different methods have been proposed for the reduction of skin friction drag such as bubble-induced skin friction drag reduction [3], drag reduction using textured hydrophobic surfaces [4] and active wall motion induced drag reduction [5]. Additionally, innovative means to reduce drag have also been developed using biomimicry [6–9].

Biomimicry may be defined as the study of design and procedural methodologies that are inspired from natural occurrences which are used to solve complex human problems [10, 11]. According to Benyus (1997) [12], biomimcry or biologically inspired designing refers to ‘a new discipline that studies nature’s best ideas and then imitates the designs and processes to solve human problems’. Thousands of years of evolution and natural selection have made natural designs, functions and processes highly optimised for survival of species such as birds, animals and insects. Bio-inspired solutions aimed at reducing drag in aircrafts and watercrafts have been created to emulate the functioning of the bodies of birds and fishes which are known to have, over time, accumulated changes in their physiology to support enhanced energy efficiency while passing through fluids. A dragonfly (*Odonata*) has triangular shaped flaps at the edges of its wings which act as vortex generators. These micro-flaps delay flow separation from the trailing edge of the wing and are highly effective in reducing drag by up to 10%, for angles of attack less than 5° as compared to a wing without any flaps. Mulligan and Rasool (2020) [13] noted that the specialised corrugated wings of a dragonfly are responsible for delaying flow separation by angles of attack up to 4° as compared to smooth wings. Other studies aimed at emulating the microstructure present on the dragonfly wing suggest a decrease in drag by about 13% on a wing with bio-inspired zig-zag edge microstructures and about 27% on a wing with zig-zag edge and pillar microstructures as compared to a flat plate wing [14–16]. Aquatic animals have been known to employ different techniques to optimise their motion through water. Shark skin is covered with several microstructures that are responsible for skin friction drag reduction [17–20]. These microstructures form a complex array of riblets which are aligned in the direction of flow of the water. A reduction in drag of up to 10% was obtained by using optimised shark inspired riblets on a surface as compared to a smooth surface [20]. These shark skin microstructures better known as denticles serve the purpose of generating vortices on the surface to prevent boundary layer seperation, similar to the vortex generators which are generally found on the wings of an aircraft [21].

Birds have also been a major source of inspiration to engineers for a very long time. The trailing edges of the feathers of a Barn owl (*Tyto alba*) have serrations, known as leading edge serrations. These serrations are responsible for improving the aerodynamic performance of wings by increasing the lift-to-drag ratio. They surpassed the lift-to-drag ratio of a wing devoid of serrations at an angle of attack of 15° in a Reynolds number (*R*_*e*_) regime of *R*_*e*_ ≥ 4 × 10^4^ [22]. Lift induced drag in aircrafts is generated due to vortex shedding at the wingtips. It has been observed that the slotted wingtips of the Harris hawk (*Parabuteo unicinctus*) are efficient in reducing the lift induced drag [23, 24]. Tucker (1995) observed that at speeds in the range of 7.3 *ms*^*−*1^ to 15 *ms*^*−*1^, there was a reduction in drag by about 10 − 20 % as compared to hawks with clipped wingtips [25]. The bio-inspired structure that is of the most importance to this paper is the herringbone riblet structures found in the feather of a bird (figure 1(a)). A periodic arrangement of herringbone riblet structures inspired from the flight feathers of a Pigeon (*Columbidae*) is capable of reducing the overall drag on a surface. Chen et al. (2014) observed a reduction in drag of 17% and 21% in planar and spatial herringbone riblet structures respectively as compared to a smooth surface [26].

**Figure 1.**
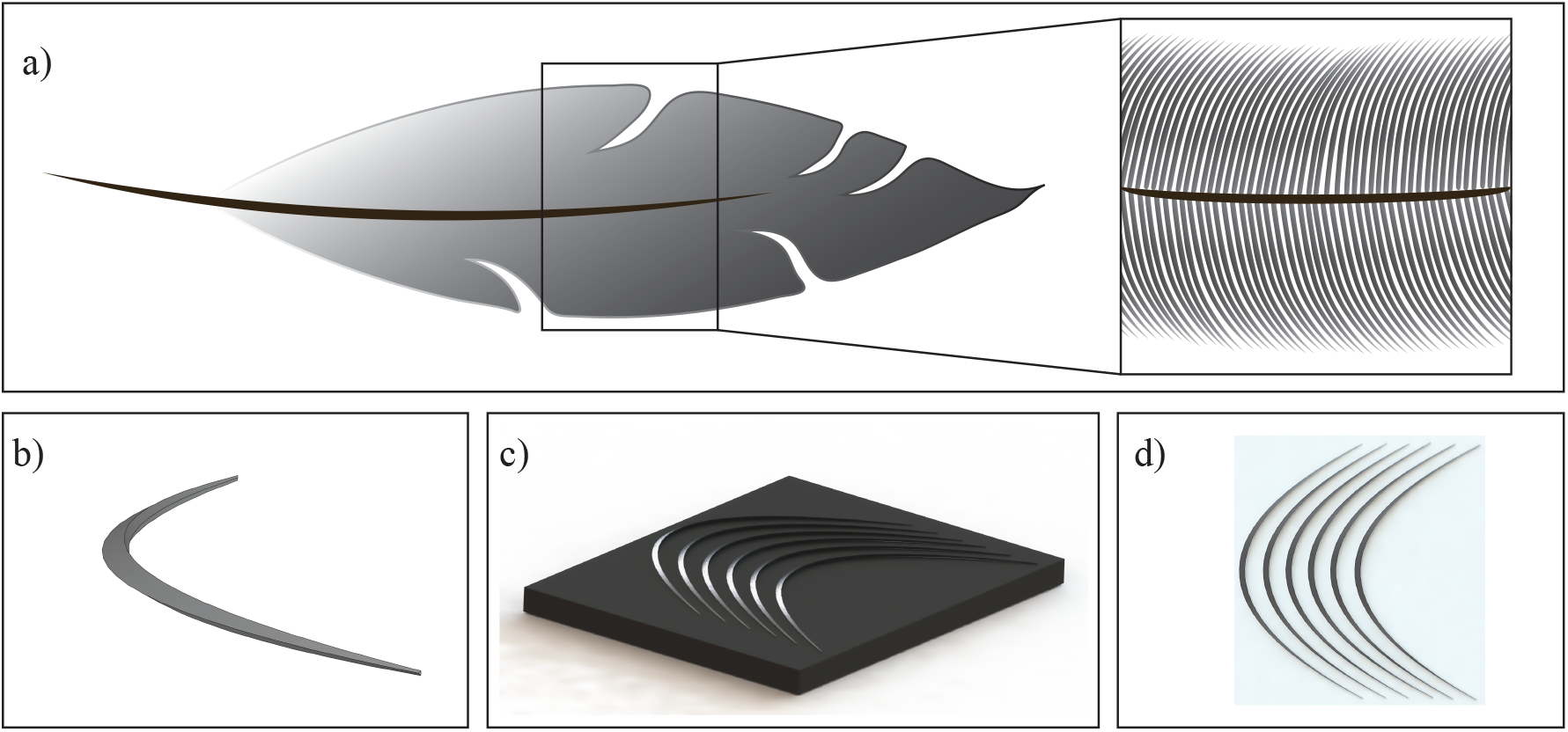
a) An image of a feather and its inherent herringbone riblet pattern. b) An individual biomimiced herringbone riblet microstructure. c) A periodic arrangement of the bio-mimiced herringbone riblet microstructures on a flat plate. d) Top view of a bio-mimiced herringbone riblet microstructure pattern on a flat plate

There have been some efforts to optimise the shape of these bio-inspired structures as well. In a periodically arranged herringbone riblet drag reduction study by Chen et al. (2014) [26], the angle between the herringbone armlets was optimised to achieve best drag reduction possible and was found to be 60°. In a similar study, Ott et al. (2020) tried to optimise the design of a shark skin denticle inspired from the Mako Shark (*Isurus oxyrinchus*) where an individual denticle structure was parameterised and optimised for overall reduction in drag by the theory of design of experiments [21]. A new and upcoming alternative to the theory of design of experiments is machine learning [27]. Machine learning in fluid dynamics has found its applications not only for design optimisation but also for analysis, control dynamics and flow estimations [28]. Machine learning can be defined as a set of processes or methods that can ‘learn’ input data and use the inherent historical patterns in the data to automate solutions of problems of complex nature [27, 29]. Machine learning in fluid mechanics has found a large variety of applications such as enhancing predictions of wake mixing in low-pressure turbines [30–32], accuracy enhancement of shock-capturing models [33] and super-resolution of turbulent flows [34]. Recently, machine learning has been used to predict the physics of flows by incorporating it in the already existing computational fluid dynamics (CFD) solutions to decrease the computational times without loosing any significant accuracy [35, 36]. Unsupervised learning algorithms such as proper orthogonal decomposition and other modal decompositions have also been used to understand the inherent structures of flows in a greater detail [37]. Machine learning in recent studies has been subjected to optimisation tasks as well [38].

To the best knowledge of the authors, there has not been a directed effort towards blending the field of fluid mechanics with biomimicry and machine learning together. Though there has been an effort to optimise bio-inspired structures, it has been done mainly using conventional and empirical methods. This includes the use of design of experiments [21], response surface methodology [21] and selectively generated parameterised models [17, 26]. These methods of design optimisation only allow for the analysis of a very small number of designs and a small parametric space. Due to this, the effects of change in certain parameters on the target variable may get neglected. By using the combination of machine learning and CFD in this paper, we try to eliminate these issues by analysing a large parametric space. This space is nearly three orders of magnitude larger than that explored in the previous studies performed for bio-inspired shark denticle and herringbone riblet structure optimisation [17, 21]. Most of the previous work to date has involved the exclusive use of physical experiments or numerical simulations. These methods have known to be tedious and time consuming if a large design space has to be explored [39]. The use of machine learning as a supplement to CFD in this study will also allow us to computationally speedup the process of exploring a large parametric space. This means there is practically no limitation to the expanse of parametric space that can explored by the proposed framework. Such a task of optimisation, if done just by using CFD, would take a very large timeframe to reach to an optimised design. Though there have been efforts to understand the flow physics of fluid over herringbone riblet patterns [40], there have only been partial efforts to optimise the herringbone riblet microstructure within a highly constrained design space [26]. The aim of this paper is to optimise an individual bio-inspired herringbone riblet microstructure (figure 1(b)) by creating a semi-automated workflow that is an infusion of bio-inspired designing, CFD and machine learning. This workflow would involve the use of random geometry generation, parametric modelling, CFD analysis and supervised learning algorithms. This framework is an intersection of three powerful and important fields and can pave a way for future commercial implementation of optimised bio-inspired designs for better aerodynamic performance and enhanced efficiency.

## 2. Methods

The complete methodology of this study, as shown in figure 2, involves the following steps: conceptualisation and parameterisation, computer aided design (CAD), random geometry generation, CFD, dataset generation and machine learning. To optimise the design of the herringbone riblet microstructure, a pilot design is created by using CAD modelling. This design is then parameterised and constraints are set on the parameters; and multiple unique designs are generated by varying these parameters randomly. This process is referred to as random geometry generation. CFD simulations are performed on these preliminary randomly generated designs to quantitatively obtain their drag characteristics. These preliminary designs along with their corresponding CFD results form the basis of the dataset that is used to generate and train a machine learning model. This machine learning model is further used as an alternative to CFD for analysing the drag characteristics of several other randomly generated herringbone riblet structures that fill up a large parametric space. The best performing structure, i.e. with the lowest drag coefficient, within this parametric space is considered to be the most optimised design.

**Figure 2.**
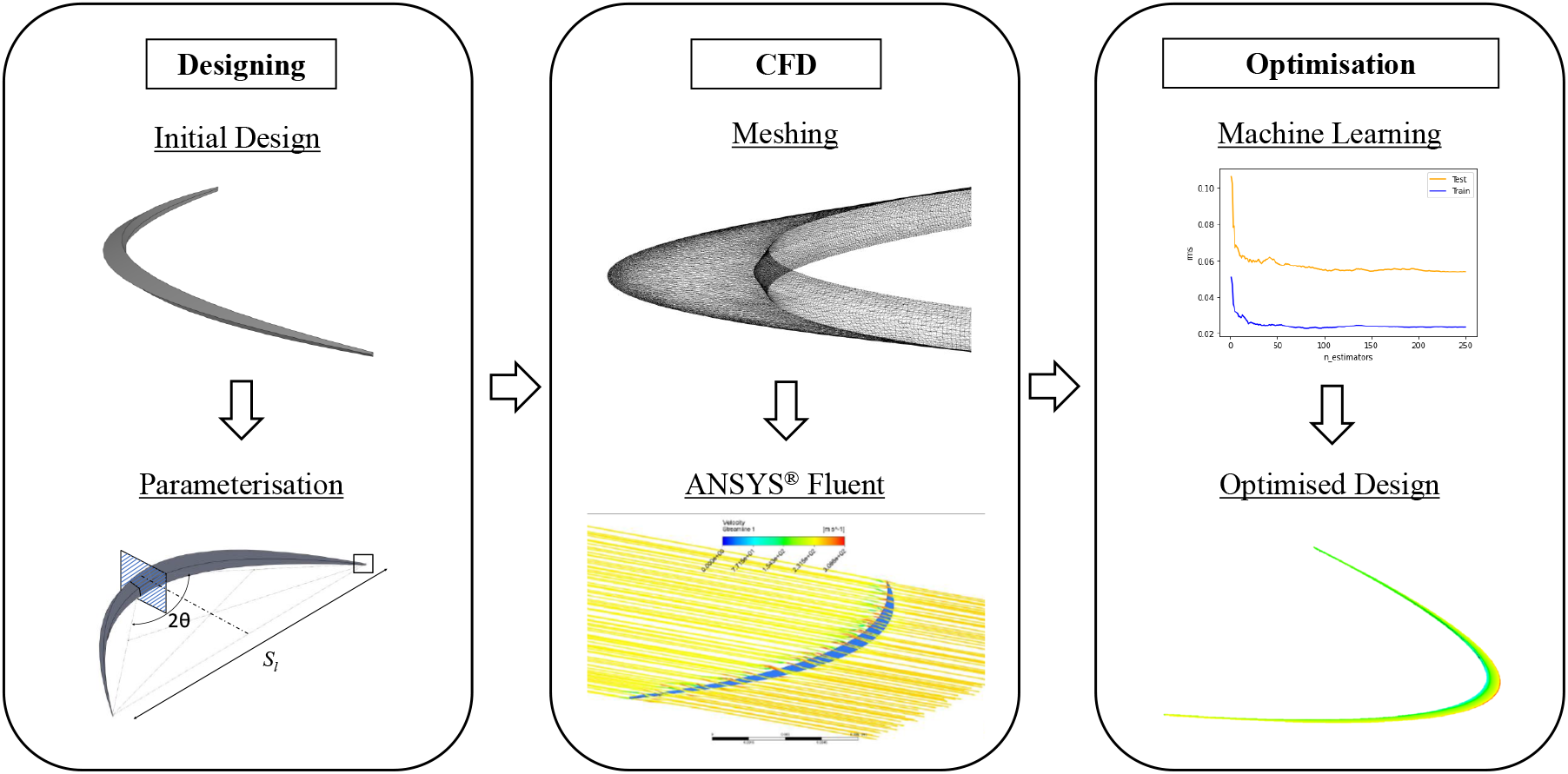
Workflow of herringbone riblet microstructure optimisation using machine learning.

### 2.1. Environment Variables

The optimisation of the herringbone riblet structure has been done to reduce the drag losses incurred by aircrafts. Though, for this study, an aircraft has been taken as the application subject, the herringbone riblet structure can also be applied to other crafts suspended in fluids. The environment variables used for the CFD analysis of the herringbone riblet structures were calculated based on the normal cruising altitude of a commercial aircraft. According to Chant (2001) [41], for a commercial aircraft the approximate cruising altitude is 11, 000 *m* and the cruising velocity is 906 *km/hr* or 250 *ms*^*−*1^. All the environment variables used are shown in table 1. As is the practice in most shark denticle and herringbone riblet optimisation studies [17, 19, 21], it can be assumed that the herringbone riblet structure is applied on a NACA0012 aerofoil. This aerofoil is symmetric and has a characteristic chord length of *L* = 68 *mm* [17]. For the ease of visualisation, figure 1(c) and figure 1(d) depict the herringbone riblet structures when applied to a flat plate. The Reynolds number (*R*_*e*_ = *UL/ν*) for this study was calculated based on the environment variables stated in table 1 with the characteristic length as the chord length of the aerofoil under consideration. Thus, the Reynolds number under consideration is approximately equal to 450, 000 which lies within the operational range of *R*_*e*_ for an aircraft (10, 000 to 1, 000, 000 [21]).

**Table 1.** Environment variables used for the study. Environment Variable Value Used

| Environment Variable | Value Used |
| --- | --- |
| Cruising altitude ($H$) | $1.100 \times 10^4 \text{ m}$ |
| Cruising velocity ($U$) | $2.500 \times 10^2 \text{ ms}^{-1}$ |
| Ambient temperature ($T$) | $2.168 \times 10^2 \text{ K}$ |
| Pressure ($P$) | $2.270 \times 10^4 \text{ Pa}$ |
| Thermal conductivity of medium ($\kappa$) <sup>a</sup> | $1.953 \times 10^{-2} \text{ Wm}^{-1}\text{K}^{-1}$ |
| Density of medium ($\rho$) <sup>a</sup> | $3.650 \times 10^{-1} \text{ kg m}^{-3}$ |
| Kinematic viscosity of medium ($\nu$) <sup>a</sup> | $3.899 \times 10^{-5} \text{ m}^2\text{s}^{-1}$ |
| Speed of sound ($a$) <sup>a</sup> | $2.952 \times 10^2 \text{ ms}^{-1}$ |

<sup>a</sup> The medium under consideration is air at an altitude of 11000 $m$ .

### 2.2. Parameterisation, CAD and Random Geometry Generation

As the first step of the process of optimisation, a design of an individual herringbone riblet is conceptualised and is translated into a digital model by using a CAD modelling software - SolidWorks^®^. This initial design CAD of the herringbone riblet microstructure has been referred to as the pilot design. For the dimensions of the pilot design, the anatomical dimensions of a feather from a Pigeon (*Columbidae*) have been used (as noted in table 2) [26]. In order to define the design of the herringbone riblet microstructure mathematically, the pilot design is parameterised. This is done by defining a set of its characteristic dimensions into variables that define the herringbone riblet structure completely. This set of parameters represents the basis for the parametric space of the microstructure. These variables, shown in figure 3, are as follows: center length (*C*_*l*_), center height (*C*_*h*_), center perpendicular length (*C*_*pl*_), flank length (*F*_*l*_), flank height (*F*_*h*_), flank perpendicular length (*F*_*pl*_), stretch length (*S*_*l*_) and half angle (*θ*), with the full angle Θ = 2*θ*. The design is considered to be symmetric in the *xy* and *xz* planes at the respective central sections (figure 3). *S*_*l*_ has been kept constant to define the dimensions of the bounding domain for the CFD portion. In order to generate a large number of designs for analysis using CFD, these defining parameters are randomly varied numerically within certain constraining bounds. These numerical constraints ensure that self intersecting and physically unmanifestable designs are not generated. The parametric constraints are also shown in table 2. Each individual set of randomly varied parameters represents an individual, randomly generated and unique herringbone riblet microstructure design. This process of random geometry generation has been employed to fill a large parametric space with several unique designs.

**Table 2.** Parameters of the herringbone riblet microstructure and their constraints.

| Parameters | Value in Pilot Design | Constraints |
| --- | --- | --- |
| Center length ($C_l$) | $300 \mu m$ | $300 \mu m - 1000 \mu m$ |
| Center height ($C_h$) | $300 \mu m$ | $100 \mu m - 300 \mu m$ |
| Center perpendicular length ($C_{pl}$) | $200 \mu m$ | $300 \mu m - 900 \mu m$ |
| Flank length ($F_l$) | $30 \mu m$ | $1 \mu m - 50 \mu m$ |
| Flank height ($F_h$) | $30 \mu m$ | $1 \mu m - 50 \mu m$ |
| Flank perpendicular length ($F_{pl}$) | $20 \mu m$ | $1 \mu m - 50 \mu m$ |
| Stretch length ($S_l$)<sup>a</sup> | $0.02 m$ | Constant $0.02 m$ |
| Half angle ($\theta$)<sup>a</sup> | $30^\circ$ | $21^\circ - 89^\circ$ |

<sup>a</sup> The value of stretch length has been kept constant in accordance to the actual length between the edges of herringbone armlets in a Pigeon feather.

**Figure 3.**
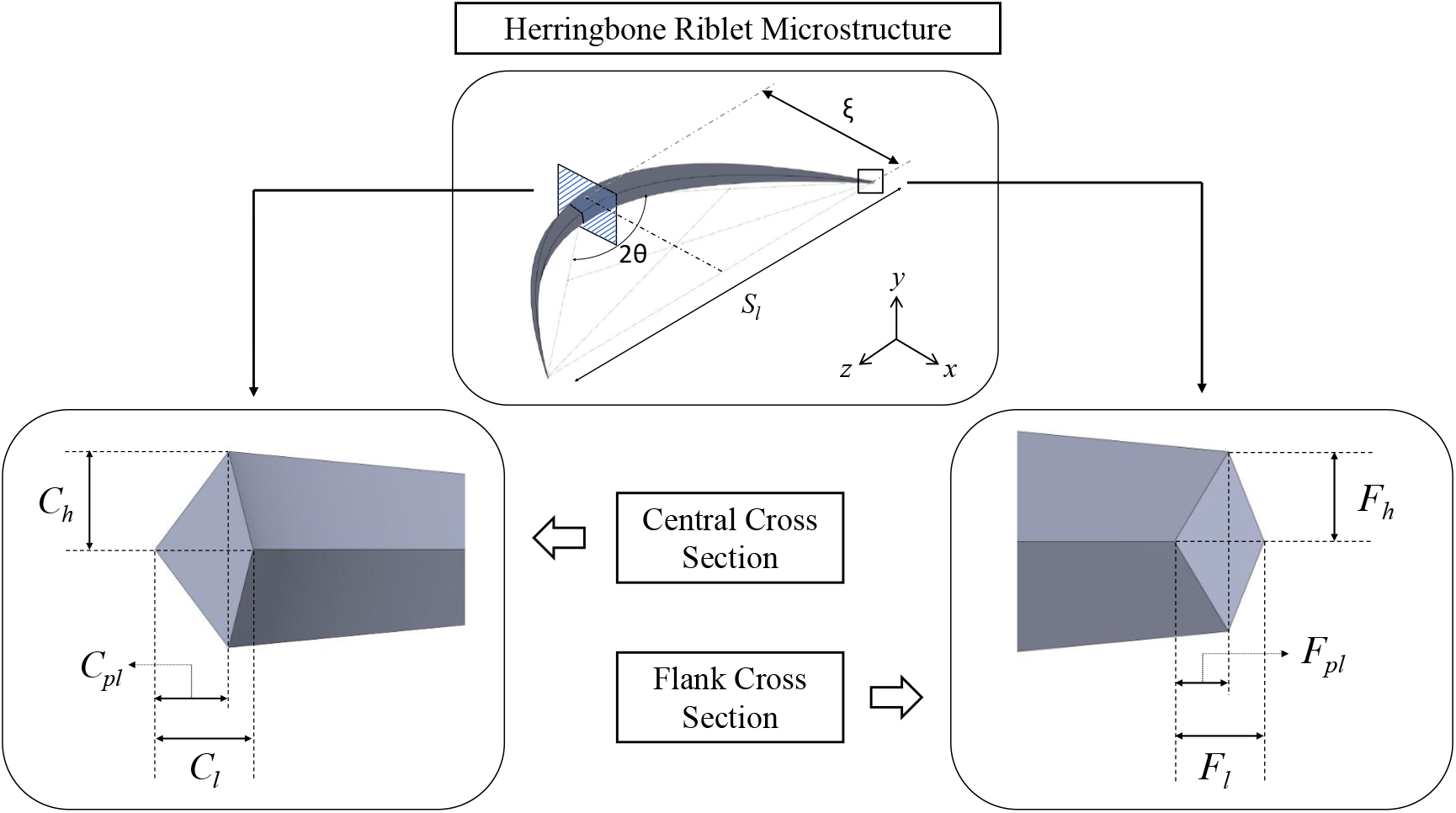
Parametric representation of a herringbone riblet microstructure.

### 2.3. Computational Fluid Dynamics

The next phase of this optimisation framework involves performing numerical simulations by using CFD on indvidual herringbone riblet structures (i.e. not placed on airfoil or flat plate). Firstly, the pilot herringbone riblet design is subjected to CFD simulations to obtain its drag characteristics. This CFD part of the optimisation process can be divided into two major categories: meshing and analysis, both of which were performed on the CFD software ANSYS^®^ Fluent v2019R2. A rectangular domain was generated such the domain size was six times the length of *S*_*l*_. A boundary layer mesh was generated near the surface of the herringbone riblet microstructure to ensure precise calculation of fluid interaction with the walls of the structure. The value of *y*^+^ was chosen to be *y*^+^ *<* 1 and the number of points in the boundary layer was set between 30 − 40 [42]. The CFD analysis of the pilot model involved using the generated mesh in the computational solver. All of the initial reference values that were used to setup this CFD solver have been noted in table 1 under the environment variables. A density based solver is used because of the high cruising velocity of 250 *ms*^*−*1^ or *M* ≃ 0.8. For the spatial discretisation of flow, second order upwind methods were used. An implicit formulation was used with a least squares cell based gradient method. The SIMPLEC algorithm was used for pressure-velocity coupling. For the calculations, a viscous *k* − *ω* shear stress transport (SST) turbulence model was employed due its ability to generate highly accurate calculations in conditions where large pressure gradients are inevitable [21]. This CFD solver solves the governing equations of fluid mechanics by using the Reynolds Averaged Navier Stokes (RANS) equations. Every herringbone geometry is tested at a zero degree angle of attack since in a local scope the flow around the body of a bird on which the herringbone riblet structures are found can be considered to be at a zero degree angle of attack [21]. In order to assess the influence of the number of mesh elements and the mesh sizes on the final calculated drag values, a convergence study was performed. The meshes analysed were varied between the number of elements in the range of 6 × 10^5^ and 1.1 × 10^6^. The variation in the drag value within these bounds was found to be 1.47 %, which is an acceptable grid convergence level [21].

## 3. Results and Discussions

This section describes the optimisation process for the herringbone microstructure. The machine learning methods and formulations are also explained. The performance of the optimised herringbone riblet microstructure is explained in terms of relevant metrics and flow physics.

A herringbone riblet microstructure is inherently symmetric and hence it must produce zero lift force at a zero degree angle of attack. Simulation results complement this assumption as most of the obtained lift force values were negligibly small (10^*−*4^*N* − 10^*−*6^*N*). Because of the large difference in the order of magnitude between the drag force and the lift force (lift force being two orders of magnitude smaller), it is safe to focus only on the drag force results and its corresponding drag coefficients (*C*_*d*_) for the objective of drag reduction [21]. The magnitude difference between the drag and lift force can be observed from the scatter plot between drag and lift coefficients (*C*_*l*_) shown in figure 4. The subsequent section will obtain and analyse the results of using these *C*_*d*_ values with machine learning to obtain an optimised design of the herringbone riblet microstructure.

**Figure 4.**
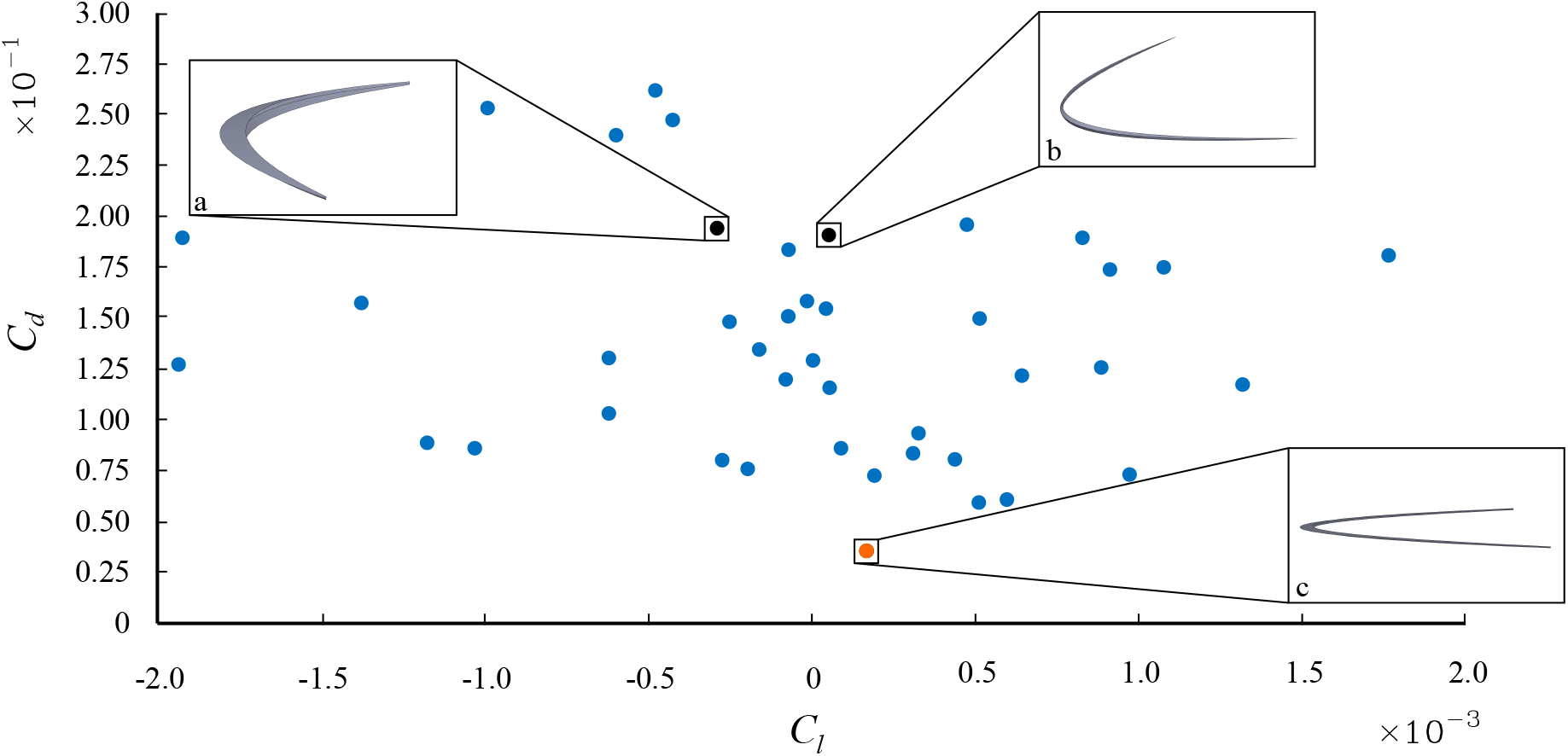
Scatter plot of *C*_*d*_ with respect to *C*_*l*_ for 41 randomly selected herringbone designs tested in CFD. a) An intermediate herringbone riblet design used in the training dataset. b) Pilot herringbone riblet design. c) Final optimised herringbone riblet design.

### 3.1. Dataset Generation

Another 107 randomly generated herringbone riblet designs were also simulated using CFD. Each of these 107 randomly generated designs derived a particular *C*_*d*_ and *C*_*l*_ value which was calculated from the drag force obtained from their CFD simulations. The formulae for the calculation of the drag coefficient and the lift coefficient are

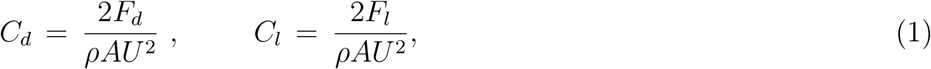

where *F*_*d*_ is the drag force, *F*_*l*_ is the lift force and *A* is the frontal area [17]. The next step of the optimisation workflow (figure 2) involves the use of machine learning to learn the relation between the herringbone riblet structure and their corresponding coefficients of drag. In order to train a machine learning model, it is mandatory to generate a mathematical dataset from the CFD results. The collection of all of the 107 randomly generated herringbone riblet structures and their corresponding *C*_*d*_ values (obtained from CFD) is the base dataset for this study. Here, each individual pair of herringbone riblet structure and its corresponding *C*_*d*_ value is known as a data point in the base dataset. Since the herringbone riblet structure parameters form the basis of this parametric space, they also correspondingly represent the feature vectors of the dataset. Also, the *C*_*d*_ values form the target variables of the dataset as they are a function of the parametric space. The base dataset used in this study is initially divided into two other other datasets. These are the training dataset (containing 86 herringbone structure - *C*_*d*_ pairs) and the testing dataset (containing 21 herringbone structure - *C*_*d*_ pairs). This corresponds to a *training* : *testing* = 80 : 20 dataset split. The machine learning models that are investigated in this study are trained on the training dataset and then tested for their performance (ability to learn) on the testing data. The best performing machine learning model is used further in the optimisation process.

### 3.2. Machine Learning

To understand the ability of different models that were investigated in this study to learn our dataset, two standard performance metrics employed for supervised learning models were used. These were the *R*^2^ *Score* and the explained variance score (*EV*). Mathematically, these metrics are defined as follows,

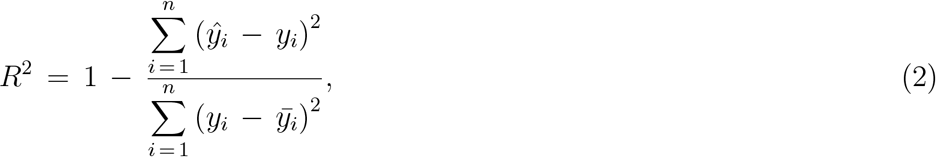

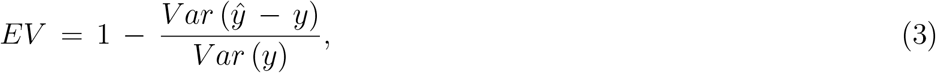

where *ŷ* depicts the predicted value, *y* represents the actual value and 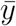 is the mean of all values. The *R*^2^ *Score* and *EV* are bounded by a maximum value of 1, which represents the best possible prediction. Initially, different regression models such as linear regressor, k nearest neighbors regressor (KNN), support vector machine regressor (SVM), decision trees, random forests and other boosting regressor methods were trained on the training data. These were then tested on for their performance on the testing data. Each of the aforementioned regressor methods used, initially resulted in a low *R*^2^ *Score*. This was attributed mainly to the large scatter in the data (figure 4) which resulted in a very low correlation between individual features. This issue of a large scatter in the data was solved by using a logarithmic transformation on the target variable.

Additionally, it was observed that dropping the *F*_*l*_, *F*_*h*_ and *F*_*pl*_ features from the dataset only had a negligible impact on the training of these models. This can be ascertained as the *R*^2^ *Scores* and explained variances (*EV*) remained constant even when these features were excluded from the dataset. These parameters physically do not alter the geometry of the herringbone riblet microstructure to a large extent as they are constrained to be almost five to ten times smaller than the other major parameters of length. This could possibly explain the constancy of the performance metrics since these parameters would marginally affect the target variable, that is the *C*_*d*_ value. Similarly, *S*_*l*_ was discarded as it is constant for every herringbone riblet design and hence does not affect the training of the machine learning model. Thus only four features were used for finding the optimised microstruture design, as the other features have a negligible impact on the final design.

#### 3.2.1. Transformed Target Regressor

The logarithmic transformation for this particular study has been performed by using the transformed target regressor. A transformed target regressor is used when there is a need to alter the target values by some sort of scaling. These scalings can be of various types including sigmoid scaling, reciprocal scaling and logarithmic scaling. For the data used in this study, it was found that the logarithmic transformation of the target values resulted in a greater *R*^2^ *Score* and a greater *EV* score as compared to normal target values for all regressors previously used. Among the regressors, the linear regression with logarithmically transformed targets gave the best results. A linear regression model tries to generate a best fitting line that describes the data. In a multi-dimensional problem such as the one presented in this study, this type of regression is also called the multiple linear regression. In such a scenario, the best fitting hyperplane is generated. The obtained *R*^2^ *Score* for this multiple linear regression model between the predicted and true values of the testing dataset was found to be 0.9644 and the obtained *EV* score was 0.9652. It can be easily observed that, *R*^2^ *Score* ≃ *EV*. This suggests that the error of the values predicted by the model is unbiased. This means that the model is largely able to capture the true relationship between the features and the target. As a measure of the average error between the predicted testing values and the actual testing target values, the mean absolute percentage error (*MAPE*) was also calculated which is,

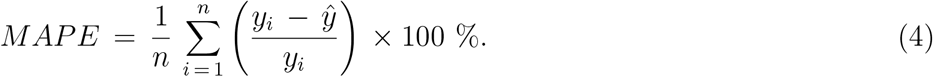

The *MAPE* is a percentage and is bounded by a lowest value of 0 which represents no error in the predictions made by the trained machine learning model. For the transformed target regressor used in this study, the *MAPE* was found to be 6.1766 %.

### 3.3. Optimisation and Drag Reduction

Once the optimal machine learning model was selected, the next step was to use it as a substitue for CFD simulations to predict the *C*_*d*_ value of several other randomly generated herringbone riblet microstructure designs that fill up a large design space. For this, an additional transformed target regressor model (as discussed in section 3.1.1) was trained with all of the 107 available data points which constitute the base dataset. Once the linear regression model with a logarithmic target transformation was trained, a mathematical relation between the target variable *C*_*d*_ and the parameters (used as features in the training dataset) was developed. This equation represents a generalised formula to obtain the *C*_*d*_ value for any given herringbone riblet geometry and is,

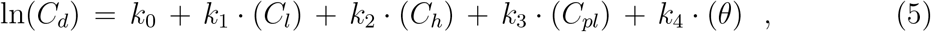

where, ln(*C*_*d*_) refers to the natural logarithmic transformation of the target variable. *k*_0_ is the target intercept and *k*_*i*_, *i* = 1, 2, 3, 4 represent the coefficients for the feature variables. The values of these constants can be obtained from the transformed target regressor model. The drag coefficient equation for any parameterised herringbone riblet is as follows,

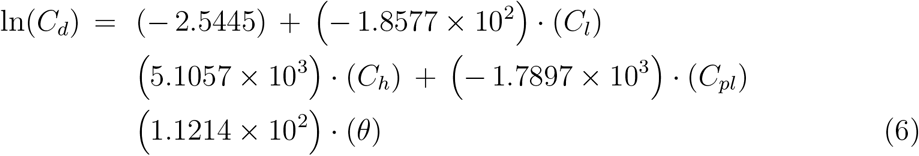

By using this trained machine learning model, the final optimised design is obtained. Firstly, 10000 randomly generated herringbone riblet structures were generated by using the methodology stated in Section 2.2. This randomly fills up the parametric space which can be further analysed for the optimisation task. The *C*_*d*_ value for each of these geometries was then predicted by using the trained machine learning model instead of calculating it with CFD. The machine learning model predicts the *C*_*d*_ value by calculating it from Equation 6 unlike the CFD simulations which calculate the drag value by solving governing equations of fluid dynamics. Hence, using a machine learning model as a substitute for CFD drastically reduced the testing time and enabled a rapid exploration of the parametric space for the optimisation process. Once the predictions of the *C*_*d*_ value for all of the 10000 randomly generated herringbone riblet designs were assimilated, the geometry corresponding to the least predicted *C*_*d*_ value amongst these was chosen as the most optimised design. Figure 5 shows three different geometries that were used to predict their *C*_*d*_ values. Figure 5(a) shows the herringbone riblet design at some intermediate *C*_*d*_ value. Figure 5(b) depicts the worst performing design while figure 5(c) shows the most optimised herringbone riblet design. The values of the defining feature variables of the pilot and optimised designs have been mentioned in table 3. This lowest predicted value of the coefficient of drag 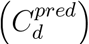 for a herringbone riblet design was observed to be 3.4103 × 10^*−*2^. To ascertain the value of *C*_*d*_ for the optimised design, the structure corresponding to this *C*_*d*_ value was tested using CFD in Ansys Fluent^®^. The true value of the coefficient of drag 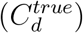 thus obtained from the CFD run is found out to be 3.5664 × 10^*−*2^. The percent error between 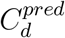 and 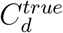 values is 4.377 %. This prediction error is even lower than the *MAPE* found for the predictions on the testing data from the transformed target regressor (section 3.2.1). This small error in the predictions suggests that the transformed target regressor model is able to learn the relation between the herringbone riblet geometries and the corresponding *C*_*d*_ values quite accurately [43].

**Table 3.** Feature Parameter Values of the Optimised Design.

| Parameters | Value in Pilot Model | Optimised Value |
| --- | --- | --- |
| Center length ($C_l$) | 300 $\mu m$ | 794 $\mu m$ |
| Center height ($C_h$) | 300 $\mu m$ | 102 $\mu m$ |
| Center perpendicular length ($C_{pl}$) | 200 $\mu m$ | 875 $\mu m$ |
| Half angle ($\theta$) | 30° | 32° |

**Figure 5.**
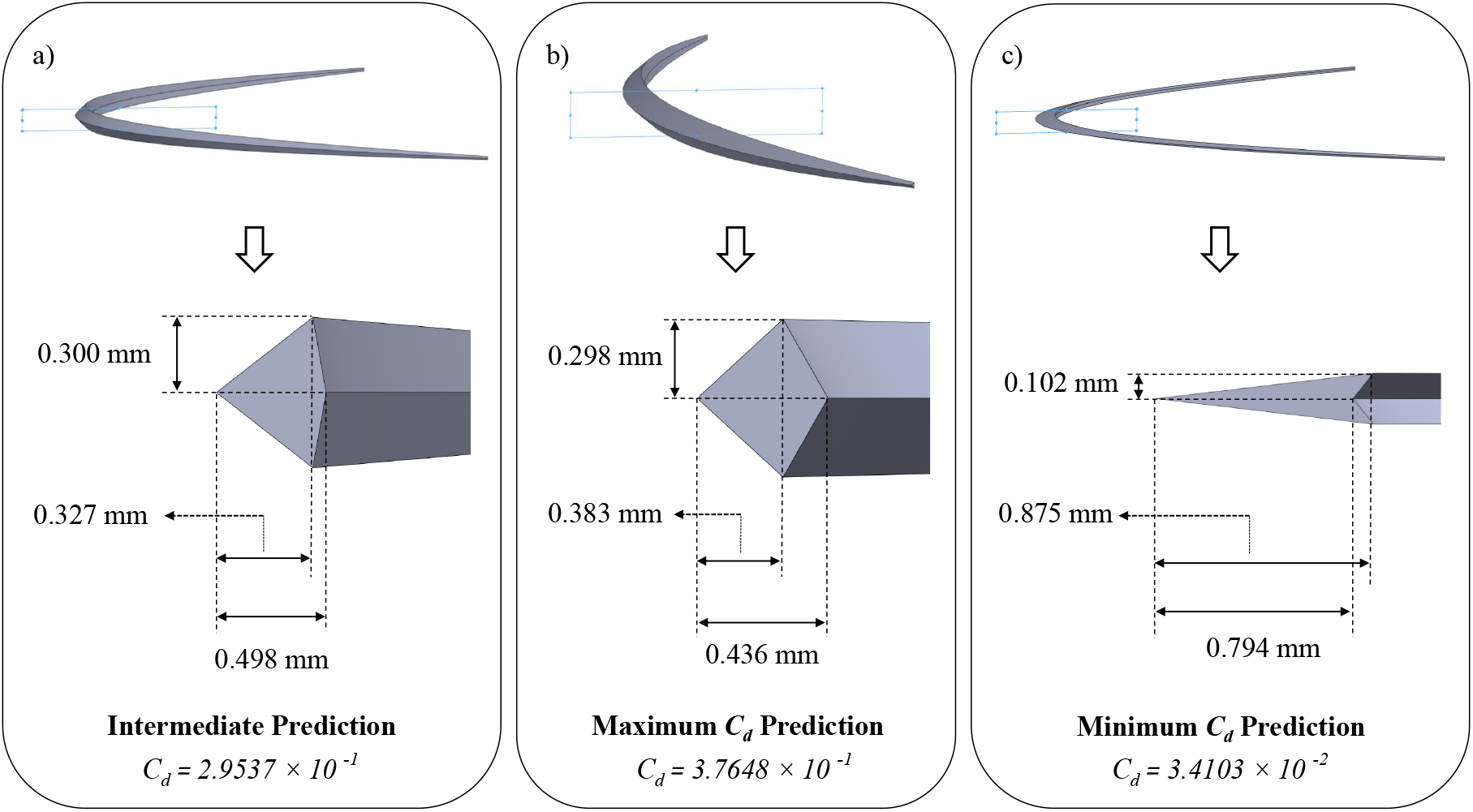
Herringbone riblet structures for (a) an intermediate design, (b) the worst design and (c) optimised design.

Chen et al. (2014) [26] used rudimentary experimental methods to find the optimum angle between the herringbone armlets. However, they only considered five different full angles between the armlets which lead to a very small design space. Though they did not optimise the structure for other parameters, they were able to experimentally conclude that the optimum half angle between the herringbone armlets is *θ* = 30°. It can be noted that an optimum half angle of *θ* = 32° was obtained in this study which is fairly close to the value obtained in previous research [26]. The drag reduction from this optimised herringbone riblet microstructure can be mainly attributed to the small separation bubble formed in the immediate wake of the structure. Figure 6(b) suggests a sharp increase in the turbulent kinetic energy in the wake of the herringbone riblet microstructure at its central cross section (figure 6(a)). The short, turbulent, separation bubble region that can be observed from the velocity streamlines in figure 6(c), may be responsible for a low pressure region in the wake of the structure which may ensure the suction of external flow towards the inner fludic regions supporting the prevention of flow separation downstream. This suggests a functioning similar to that of a shark skin micro-denticle which is known to prevent flow separation on the skin surface and is known to decrease the overall drag by benefiting from a separation bubble in the wake of the structure [17]. The intense curvature in the morphology of the armlets of the herringbone riblet microstructure are also responsible for the production of a turbulent flow in the form of vortices in the immediate wake of the structure. It has previously been shown that vortices tend to alter the size of the separation region in the fluid flow from its trailing geometry by reducing pressure and by transporting the higher energy main stream flow inwards by energising the boundary layer [44]. The vortices generated by the armlets of the herringbone riblet microstructure are also responsible for energising the boundary layer and for preventing boundary layer flow separation [17, 45]. Chen et al. (2014) [26] also suggests that these vortices generated by the herringbone riblet microstructure can reduce the fluidic drag by preventing a chaotic flow beyond the viscous sublayer or the boundary layer. Overall, it can be observed that the herringbone riblet microstructures function like a vortex generator which has been proven to promote a reduction in drag in various studies [45–47].

**Figure 6.**
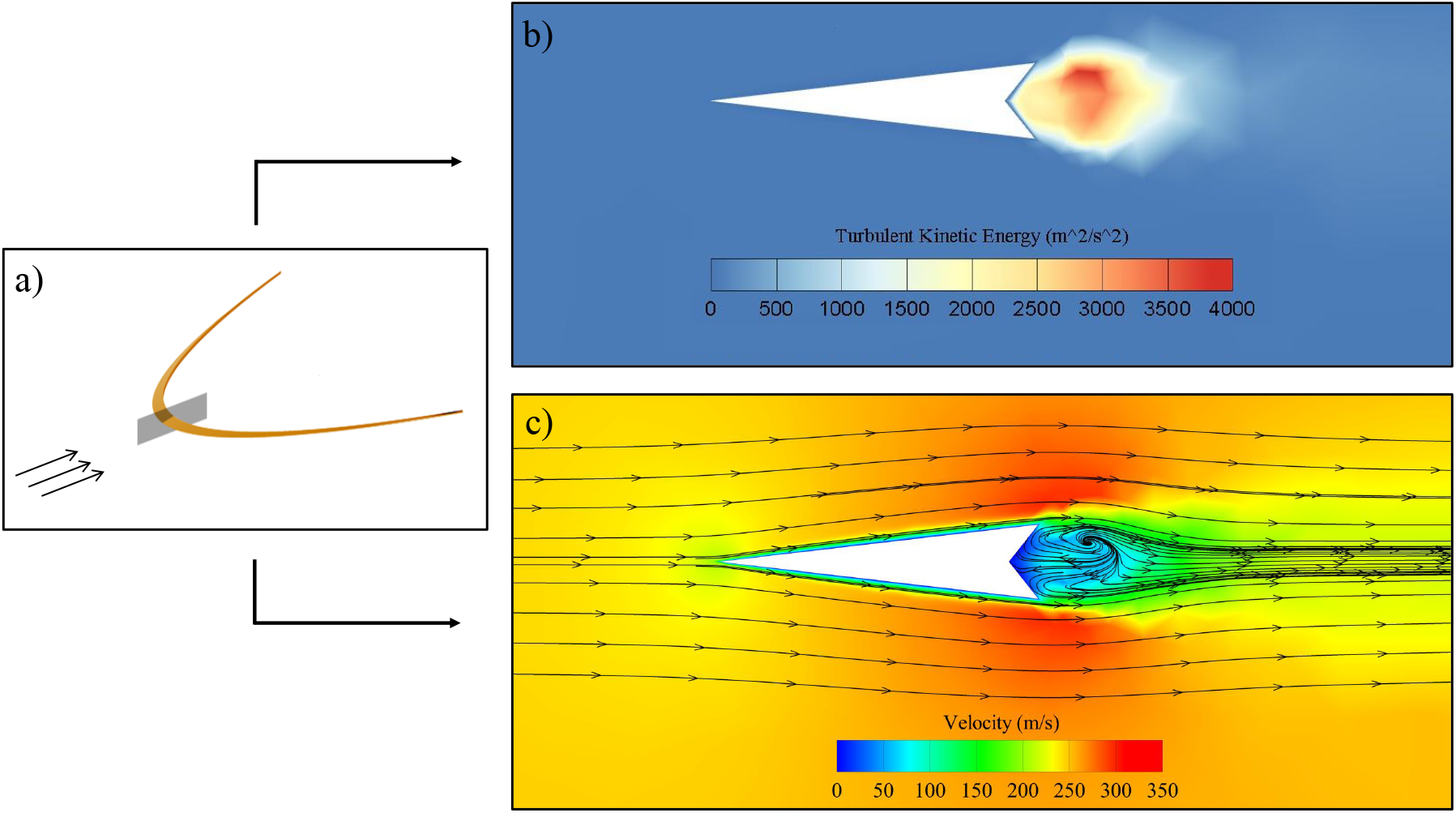
a) The central cross sectional plane of the optimised herringbone riblet design. The arrows depict the direction of flow. b) Contours of turbulent kinetic energy and c) velocity streamlines superimposed on velocity contour in the wake of the optimised herringbone riblet microstructure across its central cross section.

## 4. Conclusions & Future Work

This study traverses through fields of biomimicry, CFD and machine learning. The designs of an individual herringbone riblet microstructure inspired from a bird’s feather were investigated for their property of reducing drag while moving through a fluid. An optimal geometry for an individual herringbone riblet microstructure was proposed. This was done using a semi-automated workflow involving the generation of multiple unique herringbone riblet designs, testing them with CFD to obtain a *C*_*d*_ value and using these generated herringbone riblet structures to train a supervised machine learning model (transformed target regressor with multiple linear regression). The transformed target regressor model generated was quite accurate in its predictions with a MAPE of just 6.1766 % on the testing data. Further, this regression model was used to explore a large design space as an alternative to the computationally expensive and time consuming CFD calculations. For this, another 10000 unique herringbone riblet designs were randomly generated. The *C*_*d*_ values corresponding to these designs were predicted by using the trained machine learning model. Amongst these predictions, the design with the least value of *C*_*d*_ was chosen to be the most optimised design. The optimised design had a 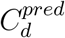 equal to = 3.4103 × 10^*−*2^. This prediction pertaining to the optimised design was highly accurate with a prediction error of just 4.377 % with respect to the *C*_*d*_ value calculated from CFD. This work demonstrates that the herringbone riblet microstructure has a functioning similar to that of a vortex generator. This study also suggests that the naturally existing evolved designs can be used to take inspiration to find solutions of engineering problems. These bio-inspired designs may not always be the best suited solutions as they are and may certainly require optimisation to extract the highest possible performance. The downfalls of some of the previous work done in this field were also addressed effectively by exploring a larger design space in a smaller time frame. This large design space was nearly three orders of magnitude larger than that explored in previous works. The optimised design obtained in this study will be able to reduce the drag encountered by crafts moving through fluids to a greater extent and should result in a greater flight efficiency.

A rigorous study involving the exact effect of such surface structures on the flight efficiency can be explored in the future by both experimental and computational approaches. Other future works may also analyse the global effects of multiple optimised herringbone riblet structures on the drag forces encountered. The relative spacing and orientation of these microstructures at different angle of attacks and in varying fluid conditions may also be analysed. Other advanced machine learning algorithms such as genetic programming and reinforcement learning may as well be used for this optimisation task in the future. The new workflow that has been devised in this study may also be used on a larger scale for the optimisation of designs in different fields of engineering.

